# Alpha oscillations control cortical gain by modulating excitatory-inhibitory background activity

**DOI:** 10.1101/185074

**Authors:** Erik J. Peterson, Bradley Voytek

**Affiliations:** University of California, San Diego, Department of Cognitive Science; Neurosciences Graduate Program; Halicioglu Data Science Institute; Institute for Neural Computation; Kavli Institute for Brain and Mind, La Jolla, CA 92093

## Abstract

The first recordings of human brain activity in 1929 revealed a striking 8-12 Hz oscillation in the visual cortex. During the intervening 90 years, these alpha oscillations have been linked to numerous physiological and cognitive processes. However, because of the vast and seemingly contradictory cognitive and physiological processes to which it has been related, the physiological function of alpha remains unclear. We identify a novel neural circuit mechanism—the modulation of both excitatory and inhibitory neurons in a *balanced* configuration—by which alpha can modulate gain. We find that this model naturally unifies the prior, highly diverse reports on alpha dynamics, while making the novel prediction that alpha rhythms have two functional roles: a sustained high-power mode that suppresses cortical gain and a weak, bursting mode that enhances gain.

## INTRODUCTION

Oscillations in the alpha (8-12 Hz) range are the most prominent feature of human visual cortical activity. Alpha is widespread, observed over the entire posterior half of human cortex. Alpha was initially viewed as a suppressive rhythm (Jasper and Penfield, 1949; Lange et al., 2013) as alpha power is maximal in visual regions when eyes are closed and the person is relaxed (Klimesch, 2012). Numerous studies over the last century suggest that this simplified characterization is incomplete because the physiology of alpha is quite complex. Alpha oscillations have been shown to rhythmically modulate population spiking (Haegens et al., 2011; Snyder et al., 2015), with alpha phase influencing information transfer (Miller and Buschman, 2013), stimulus detection (Mathewson et al., 2009), illusory perception (Minami and Amano, 2017; Sokoliuk and Vanrullen, 2013), and age-related working memory decline (Tran et al., 2016). The magnitude, or power, of alpha rapidly tracks the allocation of attention and memory in space and time (Foster et al., 2016; S. Palva and J. M. Palva, 2007; Samaha and Postle, 2015; Voytek et al., 2017) and influences trial-by-trial perception (Mathewson et al., 2009; Vanrullen and Koch, 2003). Individuals can even modulate their alpha frequency to optimize behavioral outcomes (Haegens et al., 2014; Mierau et al., 2017; Samaha et al., 2015; Samaha and Postle, 2015), possibly via thalamic modulation of alpha phase biasing cortical activity (Quax et al., 2017). These results suggest an active role for alpha where it serves a gating function controlled by top-down processes (S. Palva and J. M. Palva, 2007). This is contrary to the classic perspective and shows that alpha may not be uniformly inhibitory but, rather, rhythmically so, such that only certain phases of the oscillation aid information transfer (Chakravarthi and Vanrullen, 2012).

Here, our central assumption is that, to understand how alpha affects cognition, we must understand and account for the cortical circuits and dynamics with which alpha interacts. Alpha rhythms are (in part) generated in the thalamus (Roux et al., 2013; Schreckenberger et al., 2004; Vijayan and Kopell, 2012). Thalamic projections into the cortex can both drive and modulate cortical activity (Abbott and Chance, 2005). These thalamic projections are divided into independent nuclei that relay visual, auditory, and somatosensory input into distinct areas of the cortex (Sherman and Guillery, 2001), as well as to higher-order nuclei that transmit input from cortex to other cortical regions (Saalmann et al., 2012). All independent thalamic nuclei appear to generate and transmit 6-14 Hz oscillations to the cortex, with many corticothalamic field models spontaneously generating such oscillations (Breakspear, 2017). Alpha generation is likely a consequence of neuronal time constants and the six-layer structure of mammalian neocortex. Thalamic input to cortex targets layers 4 and 6, and cortical output occurs predominantly through layer 5 (Sherman and Guillery, 2001). The connectivity between these layers, and the intervening layers (2/3), is thought to be consistent across cortex. That is, while firing dynamics and fine computational and cognitive properties are highly regional (Kanwisher, 2010; Marcus et al., 2014), the underlying cortical connectivity patterns are conserved (Douglas and Martin, 2010).

The consistent formation and projection of alpha rhythms from the thalamus, combined with the broad consistency of cortical architecture, argues strongly for a common (biological) function for thalamocortical alpha. However there exists no such general account. Following from previous phenomenological work linking attention and working memory to gain modulation (Carandini and Heeger, 2011; Fries et al., 2001; Reynolds and Heeger, 2009; Saalmann and Kastner, 2009), and given the fact that alpha is associated with alterations in cortical gain or inhibition (Jensen and Mazaheri, 2010), we formalized thalamocortical alpha as physiological gain modulation (defined as a multiplicative change in neural excitability). We then developed a novel computational model of alpha where 10 Hz oscillations rhythmically modulate both excitatory and inhibitory (EI) firing. Thus, to understand alpha’s role in visual cognition we join two well-established models: one largely phenomenological and macroscopic—alpha oscillations control gain—and one mechanistic—EI inputs affect single unit gain.

This offers a unifying functional theory for human cortical alpha oscillations. We explore the validity of this across several computational models of visual cognitive tasks, and show that our current formalization of alpha as a cortical gain control mechanism accounts for the disparate range of observed alpha functions to date. Critically, the model also generates novel predictions for the time-varying properties of alpha that we confirm in both human intracranial electrocorticography (ECoG) and scalp electroencephalography (EEG) recordings.

## RESULTS

### Alpha modulation in single neurons

We measured the gain of a single leaky integrate and fire (LIF) neuron (Fig. 1a) subjected to either constant activity or a 10 Hz oscillatory modulation (Fig. 1b). Specifically, we modeled oscillations as periodic suppression of background activity. While the *net* effect of this oscillatory modulation is an overall increase in gain (Fig. 1c), there are time-varying phase effects such that gain reaches its maximum in the trough of the oscillation and its minimum at the peak (Fig. 1d). This observation is consistent with prior reports regarding the effect of alpha on neural gain (Coon et al., 2016; Haegens et al., 2011; Iemi et al., 2017). Although alpha oscillations are often treated as inhibitory, we find that oscillations *driven purely by excitatory inputs* produce an inverse pattern where gain is maximum near the peak of the oscillation and minimum near the trough, recapitulating previous observations of the effect of excitatory oscillations (Schaefer et al., 2006). Perhaps unsurprisingly, these results suggest that the physiological origin of alpha—generated either through pulsed excitatory inputs or periodic suppression—influences its mechanistic effects on the local cortical circuit.

**Figure 1.**
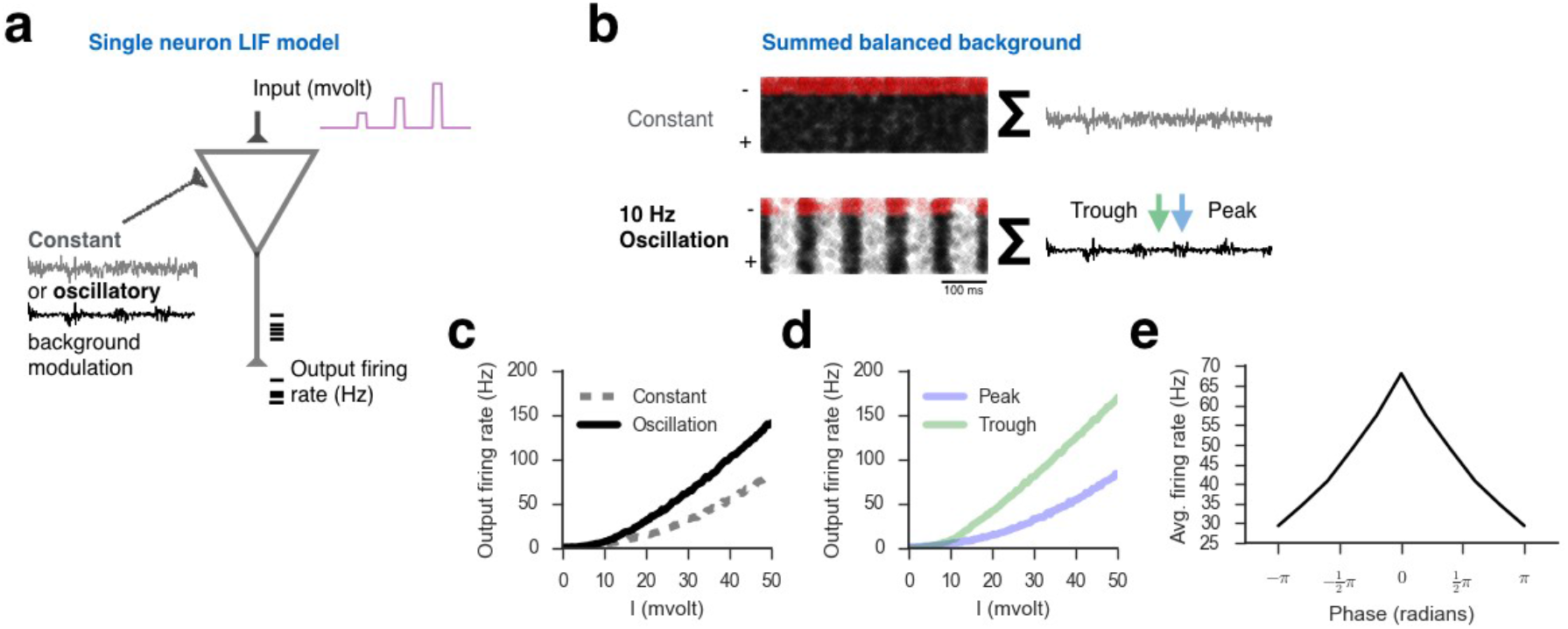
Oscillatory modulation of single unit activity. **a.** Diagram of a model LIF neuron subject to either asynchronous (constant) or oscillatory background modulation with pulses of excitatory input (purple). **b.** Spike raster of background noise *(left)* compared to the summed activity *(right)* of these balanced excitatory *(black)* and inhibitory *(red)* networks. In comparing constant *(top)* and oscillatory *(bottom*) firing modes, note how the standard deviation of the membrane potential is constant in the asynchronous case and fluctuates under oscillation (green versus blue arrows, which represent the trough and the peak, respectively). **c.** Firing rate versus input (F-I) curve of the model neuron in a under either constant *(dashed*) or oscillatory *(solid*) background modulation showing that an oscillatory background increases gain compared to a constant background. **d.** F-I curve at the peak and trough of the 10 Hz oscillation in the EI-balanced state. **e.** Firing rate as a function of oscillatory phase showing that firing is maximal at the alpha trough *(±π* rad) and minimal at its peak (0 rad).

### Alpha modulation in neural populations

To understand how oscillatory modulation affects populations of neurons, we constructed a multi-scale rate model of layer 4 of early visual cortex. This model featured a recurrently and reciprocally coupled set of excitatory and inhibitory populations, and two types of input: a “driver” stimulus, and a “modulatory” connection (Abbott and Chance, 2005). The (thalamic) driver inputs represent incoming sense information from the lateral geniculate nucleus (Fig. 2a). The modulatory inputs represent a background noise process, arising from comparatively weak thalamic connection (possibly to L6 (Olsen et al., 2012)) (Fig. 2a). We then examined modeled visual responses for two types of prototypical visual stimuli: 1) Visual Detection—a 12 ms faint burst of activity, similar to that used in many visual target detection tasks (Mathewson et al., 2009)(Fig. 2b-d), or; 2) Naturalistic Viewing—a continuous stream of noisy activity natural visual inputs (Fig. 2e-l).

### Visual detection

To study the effect of visual attention on alpha, researchers frequently use visual target detection designs wherein a faint light is briefly flashed on screen. After psychophysical thresholding to 50-70% performance, participants then must indicate whether they detected a flash on each trial and, in some designs, indicate the spatial location of the flash (Mathewson et al., 2009; Samaha and Postle, 2015). Similar to human behavioral experiments, trials are sorted by their alpha phase or power at the time of stimulus onset, and performance is examined *post hoc* as a function of alpha at the beginning of each trial.

We modeled this by presenting a 12 ms “flash” of input into our simplified model of early visual cortex. Here the flash was not a light *per se*, but an impulse of current into the model. We then examined the detection of this stimulus as a function of alpha power and phase (Fig. 2b-e). The magnitude of the stimulus was tuned until the model achieved approximately 70% detection performance *without* oscillatory modulation (see *Methods*). In line with the literature on alpha suppression (Worden et al., 2000), increasing alpha power led to decreased detection performance (Fig. 2b). However when stimulus presentation was locked to the trough of the oscillations, performance improved compared to a constant, non-oscillatory, baseline (Fig. 2c). This is expected given the model’s effect on firing (Fig. 1e). When the phase of a relatively weak oscillation (relative power = 1.5; Fig. 2b) was randomized between trials, detection performance was also improved, albeit with an increase in detection variance (Fig. 2c). These results suggest that strong, sustained oscillations suppress gain, while the lower levels of alpha typically present in visual areas during task performance (Fig. 2c,d) may represent more than a release of suppression, but may play an active, facilitatory role in visual cognition.

**Figure 2.**
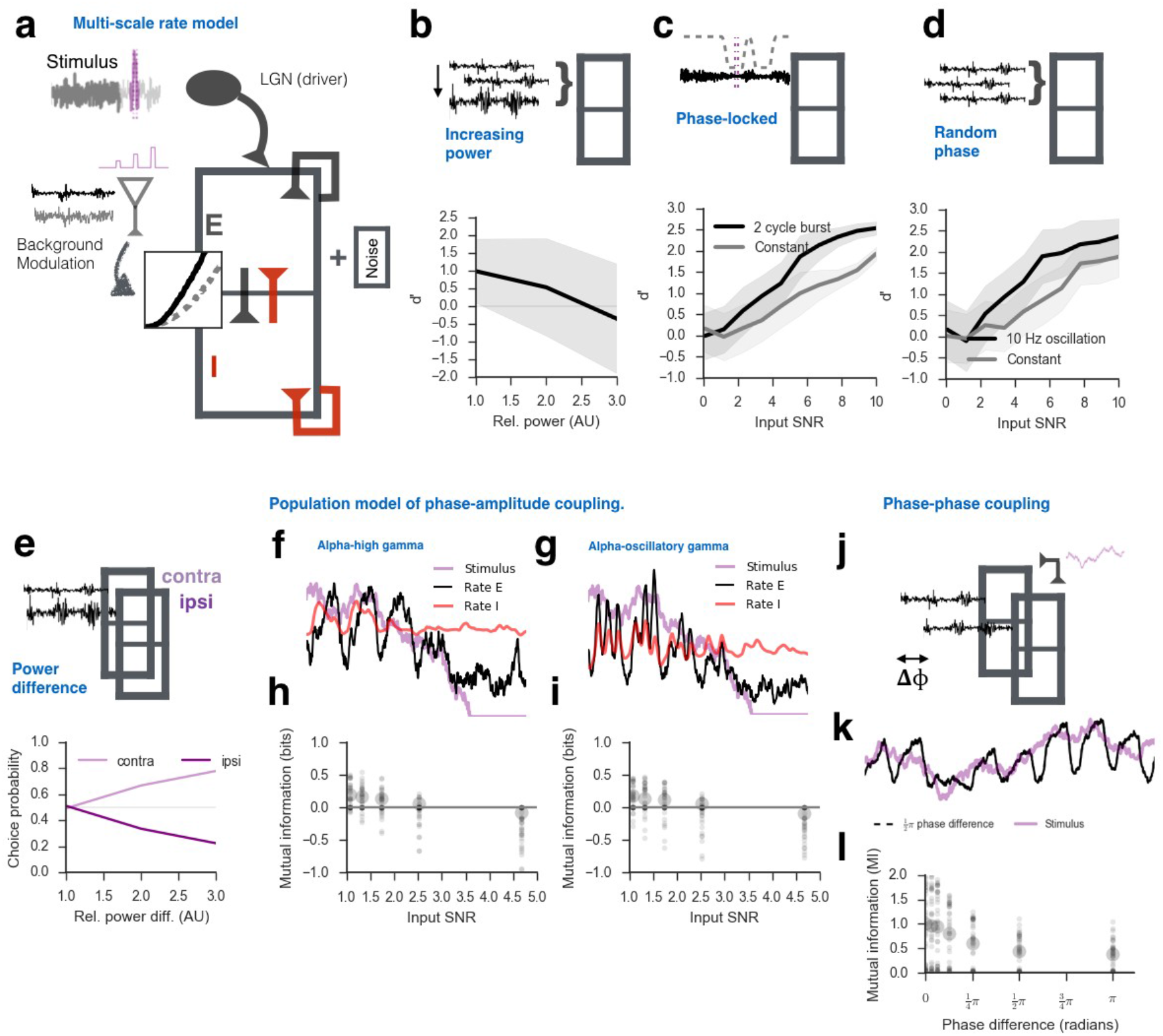
Oscillatory modulation of populations of neurons recapitulates effects of alpha power, phase, and coupling on visual task performance. **a.** Diagram of our multi-scale population mass model. Center boxes represent layer 4, the cortical input layer, and the oval the thalamic sensory input. Input here is a brief (12 ms) pulse (*violet*), simulating a rapid light flash. The dashed purple line represents the input integration window used to estimate *d’*. Background modulation is estimated in a single neuron model whose F-I curve serves to estimate population gain, following a smoothing procedure. The inset panel represents this F-I curve under constant (*grey*) or oscillatory (*black*) modulation. **b.** Simulated visual target detection (*d’*) decreased with increasing alpha power. (*Black line*, mean *d’*; *light gray*, standard error.) **c.** Phase-locked stimulus presentation showing that detection increases as a function of increasing signal-to-noise ratio (SNR), and that this effect is strong in a bursting, as opposed to constant, oscillatory state. **d.** Visual target detection also improves as a function of SNR when phase is randomized between trials, and is also better when alpha is bursting. **e.** A hemispheric gating model showing that choice probability improves for simulated lateralized visual stimuli when the alpha power difference is greatest between the two simulated visual cortical hemispheric populations. **f-g**. Example rate model time-courses under naturalistic viewing conditions. Fluctuations in alpha power lead to periodic changes in gain; such changes spontaneously generate phase-amplitude coupling in the populations in **a. h-i.** Mutual information between the stimulus and excitatory population increases with increasing white noise and type of phase-amplitude coupling (asynchronous high gamma versus oscillatory gamma-to-alpha). Small dots represent individual trials (*N* = 36); large dots represent average MI. **j.** Schematic of two independent EI populations receiving common naturalistic input, where alpha phase synchronization drifts between the two populations. **K.** Example time-courses of model output from **j. l.** Mutual information between excitatory populations decreases as the phase difference between populations increases, showing that inter-regional oscillatory phase coherence may improve information transfer.

As noted above, during visual cognitive tasks participants are frequently asked to indicate where a stimulus appears (Foster et al., 2016; Samaha and Postle, 2015; Tran et al., 2016). Alpha power is widely reported to decrease more in the visual cortical hemisphere contralateral to stimulus presentation. We therefore implemented a “hemispheric” version of our population model. Two population models were weakly coupled to produce a bilateral model of the early visual system (see *Methods*). The model was tasked with identifying on which side a visual stimulus appeared. In this model, differences in alpha power could effectively control choice probability. Decreasing contralateral, relative to ipsilateral, alpha power suppressed false positives in that hemisphere, which increased correct detection events for contralateral targets (Fig. 2e).

### Communication though coherence

In human and non-human animals, oscillatory coupling between cortical layers, and between brain regions, is believed to control information flow. Two modes of coupling are commonly reported: phase-amplitude coupling (PAC)(Canolty and Knight, 2010) and coherence (phase-phase coupling)(Fries, 2005). We studied the effect of coupling on information flow under simulated naturalistic viewing conditions.

The periodic fluctuations in gain intrinsic to our model produce two modes of PAC: slow oscillation phase coupled with asynchronous firing (*i.e.*, “high gamma”, 80-150 Hz) and the coupling between a slower alpha oscillation and a faster oscillation in the beta/gamma range (20-50 Hz; Fig. 2f-g). Note that distinguishing between these two forms of PAC is critical, as they likely have different physiological interpretations (Cole et al., 2017; Cole and Voytek, 2017; Vaz et al., 2017). In line with previous reports, both modes of oscillatory coupling lead to an increase in information flow compared to the non-oscillatory control condition (Peterson and Voytek, 2015)(Fig. 2h-i). Additionally, when the “hemispheric” model (see above) was exposed to common naturalistic input, but the phase of the alpha modulation was allowed to drift between regions such that they could be in phase or drift out of phase (Fig. 2j,k), information flow decreased as the two regions moved out of phase (Fig. 2l). This observation recapitulates theories on neural communication through phase coherence (Fries, 2005), where inter-regional phase coherency has been previously linked to maximized information flow (Buehlmann and Deco, 2010; Kirst et al., 2016), possibly via PAC (Voytek et al., 2015; Voytek and Knight, 2015).

### Contradictory effects of alpha power

Increased oscillatory power can arise from several different sources, including either increasing the amplitude of each cycle, or simply increasing the number of cycles over the temporal integration window (Jones, 2016). To simulate the former, we show that increasing the peak firing rate of a neuronal population generating oscillatory alpha increases alpha power, but decreases neuronal gain (Fig. 3a-c). In contrast, while increasing the number of oscillation cycles also increases alpha power (Fig. 3d,e) this “bursting” mode has the seemingly contradictory effect of *increasing* gain (Fig. 3f).

**Figure 3.**
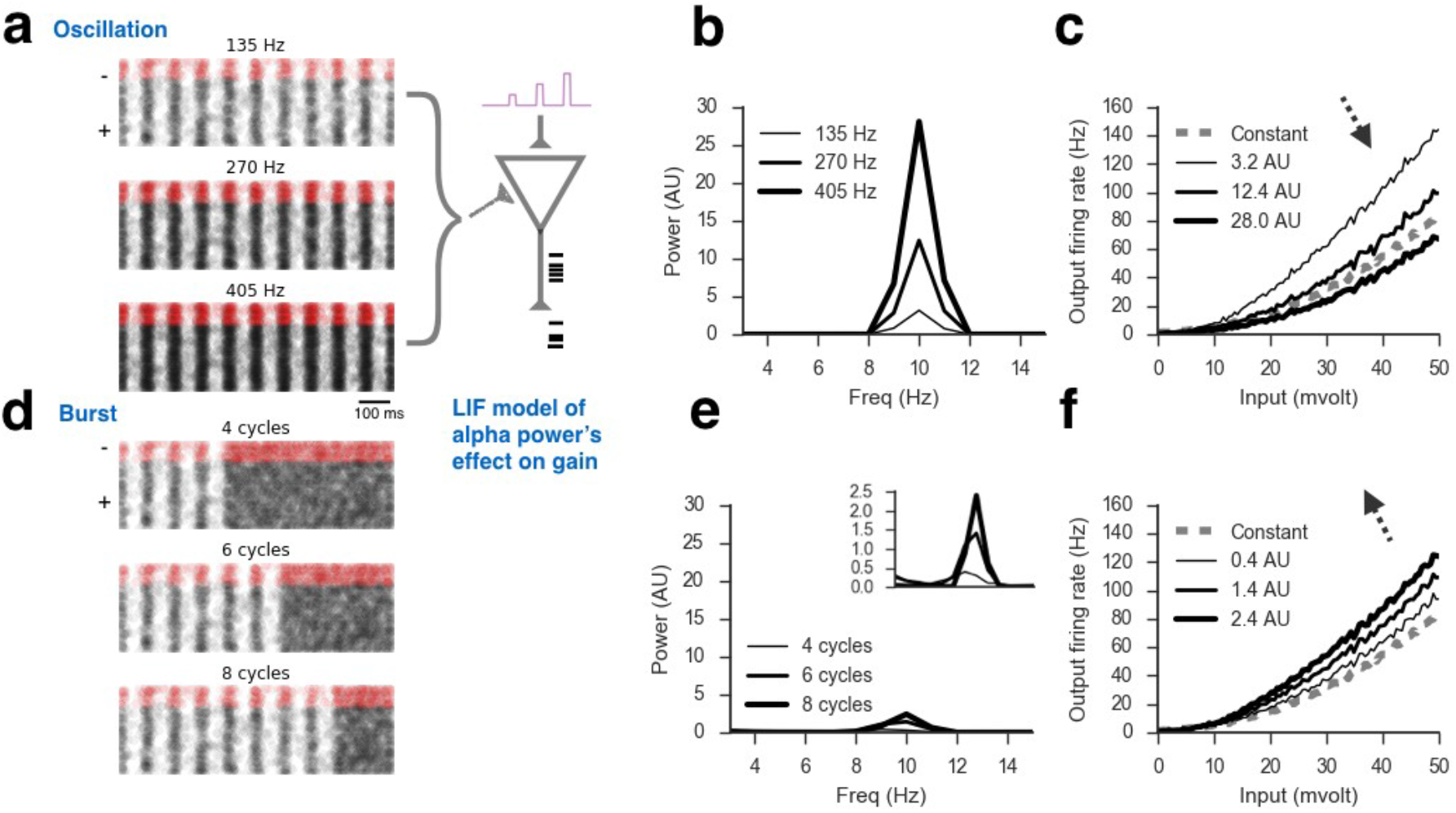
Alpha power increases can imply both increases or decreases in neural gain depending on their physiological origin. **a-b.** Alpha power increases as the peak neuronal firing rate in a sustained oscillation increases. **c.** Increases in sustained alpha power lead to decreases in F-I curve slope, indicating a decrease in gain. **d-e.** Increasing the number of alpha cycles in a short burst of oscillatory activity also increases alpha power. f. Increased alpha power due to an increase in burst number, however, increases gain (F-I slope).

That is, our model suggests that alpha power is not a singular physiological phenomenon, but rather may have two distinct functional modes: A strong, sustained mode (>5-10 cycles) that serves to suppress neuronal activity, and a weaker “bursting” mode (of 1-5 cycles) for fast, precisely-timed gain increases. Given this, we hypothesize that the sustained mode should therefore decrease the average firing rate, as well as the variance of firing, in an oscillating region. That is, sustained alpha suppresses, but stabilizes, firing. We further hypothesize that alpha bursts act in an opposite manner—to rapidly but temporarily increase gain, which is supported by several recent reports that brief (100 ms) cortical beta (13-30 Hz) bursts of just a few cycles aid in behavioral outcomes (Feingold et al., 2015; Jones, 2016). That is, our model suggests oscillatory bursting should increase both average firing and firing variance.

The above modeling results offer novel predictions regarding the physiological and functional role of sustained versus bursting alpha. To experimentally test these hypotheses, we analyzed both invasive and non-invasive human visual cortical electrophysiology while participants performed visual attention tasks.

Analysis of visual cortical EEG data shows that, during eyes-closed rest, there is a bimodal distribution of alpha power in visual areas. However, when participants are performing a visual attention task, only lower power mode is apparent (Fig. 4d). In terms of oscillation duration, task-related alpha has a median length of 0.256, corresponding to roughly 2-3 cycles (Fig. 4e). This represents a 57% reduction in length compared to rest periods (median length of 0.404 seconds; Fig. 4e). Consistent with our EEG results, the median duration of task-related alpha oscillations in our analysis of ECoG data (discussed below) was 0.285 seconds.

**Figure 4.**
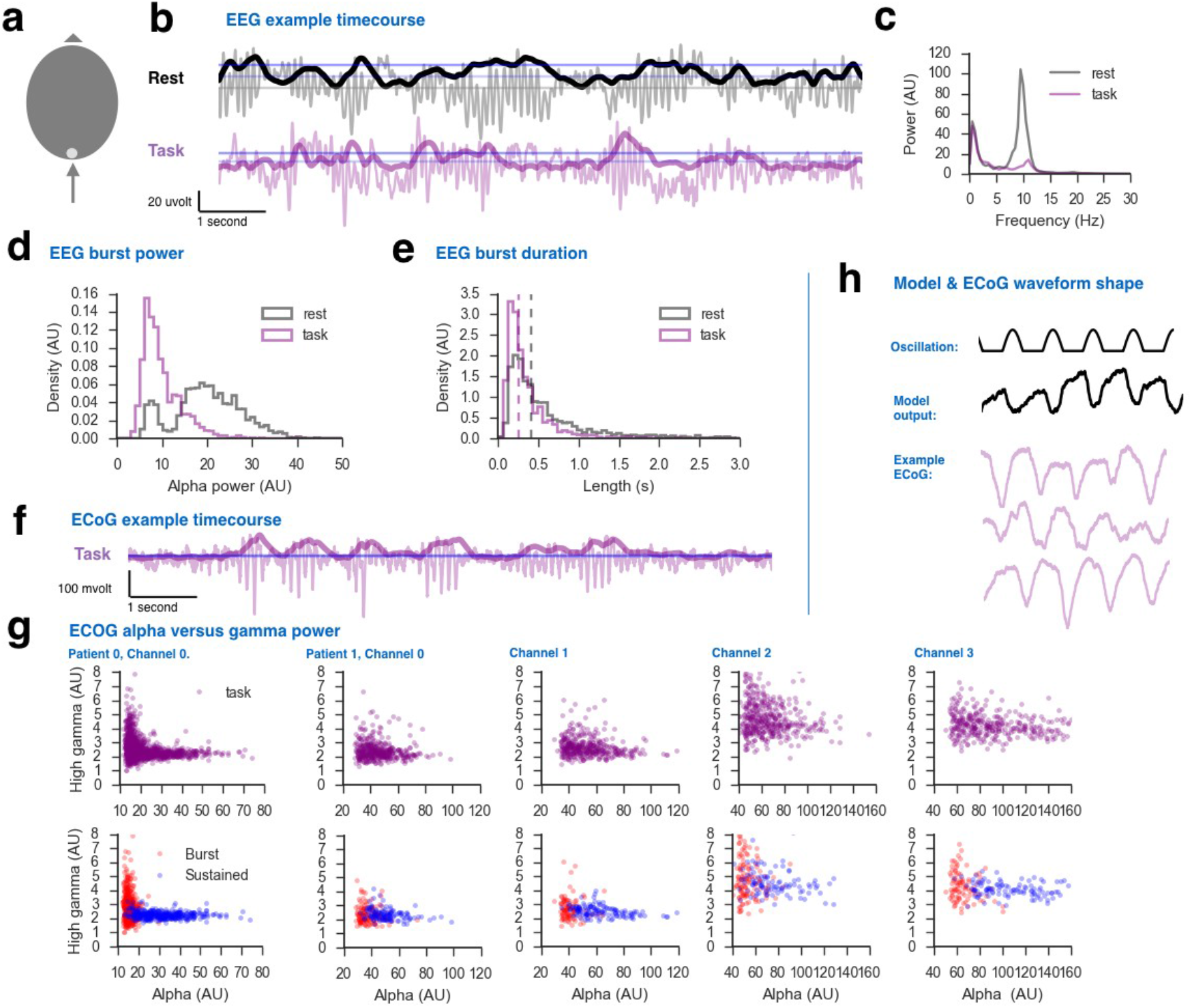
Sustained versus bursting: evidence for two modes of alpha activity in humans. **a.** All electrophysiological data was collected from posterior occipital areas (ECoG or electrode OZ in EEG). **b.** Example EEG data from both task and rest periods. Dark lines represent instantaneous alpha power, light lines represent the unfiltered electrophysiological recordings. Blue lines denote the criterion for burst onset (*dark*) and offset (*light*) criterion. **c.** Power spectra of EEG data showing significant alpha power reduction from rest to task. **d-e.** Distribution of oscillatory power (**d**) and duration (**e**) for all detection periods of oscillatory activity. **f.** Example of ECoG data (only task data was available for these participants, not rest). Dark lines represent instantaneous alpha power, light lines represent the unfiltered electrophysiological recordings. Blue lines denote the criterion for burst onset (*dark*) and offset (*light*) criterion. **g.** Relationship between alpha (8-12 Hz) and high gamma (80-150 Hz) power for two participants, both overall (*top*) and as function of oscillation type (*bottom*). Oscillations were periods of activity lasting greater than 0.5 seconds (5 cycles), while “bursts” were periods of oscillation lasting less than 0.3 seconds (3 cycles). **h.** Comparison of oscillation waveform shape between model and ECoG data.

To assess the physiological link between alpha and spiking in humans, we examined alpha as a function of asynchronous high gamma power (50-150 Hz), which is believed to index increases in multi-unit spiking activity (Manning et al., 2009; Mukamel et al., 2005). However, because of the difficulty in measuring high gamma non-invasively, due to nonneural noise sources in scalp EEG (Voytek et al., 2010b), we analyzed invasive human ECoG data from 2 patients performing a visual attention task. Specifically, we compared the relationship between high gamma and alpha power (Fig. 4g, *top panels*), classifying periods of alpha oscillation into two categories: *sustained* oscillatory alpha (> 5 cycles) and alpha *bursts* (< 3 cycles, see *Methods*). Sustained alpha was related to significantly reduced high gamma power, consistent with previous ECoG reports (Coon et al., 2016; Crone et al., 1998; de Pesters et al., 2016). In contrast, bursts showed an approximately 2-4-fold increase in gamma power and variance (Fig. 4g, *top panels*). In fact, separating sustained versus bursting oscillations explains the nonlinear correlation observed between alpha and high gamma power (or spiking)(Snyder et al., 2015)(Fig. 4g, compare *top* and *bottom* panels).

### Waveform shape

The background oscillatory process in our model is a periodic burst of background firing followed by a return to a longer period of below-baseline activity (Fig. 4h, *top*). This pattern is inspired by the highly synchronous bursts of activity typically reported in the thalamus, but is quite dissimilar to the waveforms typically observed in EEG (Fig. 4b) and ECoG recordings (Fig. 4f,g, *bottom*). When this “thalamic” pattern is used to control gain, excitatory population waveforms are remarkably similar to the patterns observed in human electrophysiology (Fig. 4g, *middle*). That is, ECoG and EEG recordings of alpha present with typically asymmetric, non-sinusoidal waveforms, featuring a comparatively narrow and sharp decreasing component and relatively broad, smooth upward profile (Fig. 4g). This is the pattern observed in our model, a pattern that is unique to balanced gain modulation. Under the same conditions, oscillatory modulation driven by excitatory input alone produces a smoother, symmetric, waveform (not shown). This is in line with new evidence that physiological information may be extracted from the waveform shape of non-sinusoidal oscillations(Cole et al., 2017; Cole and Voytek, 2017).

## DISCUSSION

We show how the diversity of changes to alpha power and phase can be understood in terms of a single mechanism—gain modulation—implemented *specifically* by periodic fluctuations in balanced EI background activity. This view of alpha unifies past theoretical views with empirical behavioral and physiological findings under a single mechanism. This is significant, as the spectrum of theories about the functions of alpha are broad, ranging from suppression, gating, sampling, and gain control to working memory maintenance and attentional allocation. A common thread to these interpretations is the modulation of information. Beginning from this common view, we began by positing a biophysical mechanism for alpha: balanced gain control(Peterson and Voytek, 2015). From there we found that our model could capture much of the existing alpha literature while offering novel predictions.

The earliest accounts treated alpha as an idling rhythm. This view is challenged by studies examining alpha as function of, for example, working memory, by showing that alpha power in posterior electrodes increases with working memory load across many paradigms (Jensen et al., 2002). The idling hypothesis was thus updated to an account where alpha acts to rhythmically inhibit neural gain (Cooper et al., 2003; Haegens et al., 2011; Iemi et al., 2017). In this updated view, regions not receiving sense information or otherwise engaging in active behavioral responses were said to be suppressed to avoid introducing noise into the larger network. When stimulated, task-relevant regions are disinhibited, restoring cortical processing.

An alternative account of alpha oscillations and working memory is the phase-coding approach, where individual neuronal spiking activity is phase-locked to the oscillation (Lisman and Idiart, 1995), offering a route to encode memory items inside single alpha cycles (Jensen and Lisman, 1998; Lisman and Jensen, 2013). While influential, both the inhibition and phase-coding models have two crucial shortcomings (Jensen et al., 2002): 1. Alpha activity occurs outside working memory and attentional tasks, and; 2. Regions implicated in working memory and attention (such as prefrontal cortex) have weak, if any, alpha.

In our model, alpha oscillations, when treated *specifically* as balanced gain modulation, can suppress visual detection. When suppression acts over long periods (*e.g.*, >1 sec) this strongly reduces the overall gain (Fig. 2b). This low-gain state reflects an “idle” state of activity, serving to inhibit (*i.e.*, reduce) firing output for any given input. Stimulus presentation during the high-gain trough of alpha maximizes detection sensitivity, giving the model a natural “phase preference” for visual stimulus detection (Fig. 1d,e and Fig. 2c). However even if phase is randomized, the average increase in gain provided by weak oscillatory activity improves detection performance compared to non-oscillatory periods (Fig. 2d). This feature offers an intuitive answer to why noticeable alpha oscillations are present even during very difficult visual tasks (Fig. 4b,f): if alpha acted *only* to suppress activity, naively one would expect oscillatory activity to cease in task-relevant regions during the most difficult cognitive processes. Yet, oscillatory activity persists.

This alpha-as-gain theory also provides for gating operations by biasing phase (Fig. 2l) and amplitude (Fig. 2e). For example, competition between hemispheres during detection of a random lateralized stimulus can be accomplished in our model by either biasing the power between two tonic oscillations (Fig. 2e, j-l), or by the focal application of “bursting” activity to the desired hemisphere (see below).

Regarding the phase-coding view of alpha, wherein asynchronous high gamma or oscillatory gamma are nested inside alpha cycles, our model generates both modes of PAC. Asynchronous high gamma PAC is thought to reflect the phasic control of the aggregate firing rate while nested beta/gamma-to-alpha PAC, in contrast, reflects the phase of a slow oscillation (alpha in this case) controlling the amplitude of a faster oscillation (20-60 Hz beta or gamma). Both types of PAC have been observed in electrophysiological recordings of human and non-human primates (Vaz et al., 2017) with visual cortex high gamma PAC preferentially coupling to alpha (Voytek et al., 2010a). However why one mode is preferred in some regions, tasks, periods, and participants is not understood. Our model suggests an answer: the transition from asynchronous high gamma to nested beta/gamma-to-alpha PAC depends largely on the connectivity of the local EI populations. That is, both types of PAC are driven by gain modulation. However whether high gamma or beta/gamma is present depends on the local connectivity. In this case the transition from high gamma to nested beta/gamma PAC follows from doubling the recurrent excitatory connection (*W_EE_*). However increasing the EI ratio (*W_IE_/W_EI_*) or the length of the inhibitory time constant (*τ_gaba_*) can also control the transition from high gamma to nested oscillatory PAC. Likewise, whether beta or gamma oscillations appear depends on the strength of local EI reciprocal connections and time constants, and the magnitude of gain changes, such that higher gain increases the relative strength of cellular input while stronger inputs favor faster oscillations (Brunel and X.-J. Wang, 2003).

A major question in neuroscience is how sense information from the periphery is represented cortically. A controversial and longstanding debate centers around whether that representation is continuous or discrete, with evidence for both perspectives. Without any special tuning our model explains both of these continuous versus discrete features. Periodic alterations in gain segment continuous sense input into, effectively, high and low informational bins. When measured experimentally, such bins may give the appearance of a “discrete” representation of the sense information. However the difference in informational content induced by the gain effect is modest when oscillatory activity is weak (Fig. 2b), as it is during task periods (Fig. 1b,f). Likewise, gain effects are strongest when noise is large (Fig. 2h,i).

### Model predictions

Though our model of alpha modulation can produce a continuous range of effects, we took a dichotomous approach in order to generate novel testable predictions. Specifically, we hypothesized that alpha oscillations could serve two distinct functional roles: 1) A strong sustained oscillation (> 5-10 cycles) for suppression, and; 2) A weaker “bursting” mode for fast, temporally-precise gain increases (of 1-3 cycles; Fig. 3). Experimentally, using both invasive and non-invasive human electrophysiology, we observe that alpha power has a bimodal distribution, consistent with our bimodal hypothesis for alpha activity. Further, only low-power, low-duration “bursts” of alpha lead to increases in high gamma power, a well-established proxy for multi-unit spiking (Manning et al., 2009; Mukamel et al., 2005) (Fig. 4g, compare *top* and *bottom* panels).

A distinction between relatively sustained versus bursting oscillatory periods is supported by recordings in thalamus of nonhuman primates, in which alpha rhythms appear in two modes: tonic oscillations and bursts of alpha power (Sherman and Guillery, 2001; 1998). The contrast between the number of bursts increasing gain while the increasing magnitude of tonic oscillations decreases gain, suggests an answer to the question of why the thalamus explores separate firing modes: bursts may serve to provide a rapid, stimulus dependent tuning of gain while tonic firing may serve to modulate or inhibit regional activity (Jensen and Mazaheri, 2010; Mazaheri, 2010). This would, as previously proposed, inhibit noisy or undesirable activity in one area from influencing other regions. Both stimulus-and internally-driven tuning, as well as homeostatic demands, may therefore seek to rapidly modulate gain. Our model suggests that fluctuations in alpha power, phase, and frequency be viewed as part of singular cortical tuning process.

Outside of normal cognitive functioning, disruptions to alpha have been implicated in numerous neurological and psychiatric disorders, most markedly in depression (Henriques and Davidson, 1991) and autism (Edgar et al., 2014). Notably, these disorders are often associated with alterations in EI balance as well as with working memory and attention impairments (Rubenstein and Merzenich, 2003; Shabel et al., 2014; Voytek and Knight, 2015). Given our current results unifying alpha and EI dynamics, it is plausible to suspect that the observed alpha and EI balance disruptions across these disorders—as well as associated cognitive deficits—reflect similar physiological mechanisms. Difficulty in measuring EI dynamics in humans has prevented exploring such links, however recent advances in estimating EI balance from the local field potential (Gao et al., 2017) may offer novel insights.

Our results show how alpha oscillations can manipulate balanced EI coupling to control gain, and suggest this control is used across a wide range of visual and cognitive operations— including attention, memory, gating, and sensory sampling. These results show that alpha is not an index of *a* cognitive process, but rather that alpha oscillations index a canonical circuit mechanism used to implement many aspects of cognition.

## EXPERIMENTAL PROCEDURES

All participants gave informed consent as approved by the appropriate IRB at the institutions where the data were collected. Modeling assumptions and methods, as well as details of data and analysis procedures, are available in Supplemental Experimental Procedures.

## SUPPLEMENTAL EXPERIMENTAL PROCEDURES

We developed a model of periodic gain control, relying on the manipulation of balanced interactions between excitatory and inhibitory neurons. Balanced here implies that increases in excitatory firing are quickly (< 3 ms) and exactly countered by an increase in inhibitory activity. With the converse being true for inhibitory activity. Modeling efforts in this work spanned two levels of analysis. First we studied how a balanced 10 Hz modulation altered the firing properties of single LIF neurons. Second we used the tuning properties of this model to develop a population firing rate model of layer 4 in early visual cortex. We then subjected this cortical model to range of visual cognition experiments to demonstrate that it can produce effects highly consistent with the existing literature. Finally we confirm the novel predictions made by our model using both invasive and non-invasive human electrophysiological data.

### Modeling assumptions

The central assumption of our model is that EI interactions within and between cortical layers are balanced, as defined above. This is supported by evidence that in awake, behaving animals, excitatory and inhibitory firing exists in a *balanced* state such that increases in excitatory firing are rapidly and exactly countered by inhibitory firing (Xue et al., 2014). The balanced state is crucial for efficient neural coding, transmission, and information gating (Peterson and Voytek, 2015; Vogels and Abbott, 2009; 2005; Womelsdorf et al., 2014) as well as for maintaining neural homeostasis and preventing seizure activity (Dehghani et al., 2016; Truccolo et al., 2014; Wendling et al., 2002). Balanced interactions can also give rise to gain modulation (Abbott and Chance, 2005; Chance et al., 2002). That is, if EI balance is maintained, but the overall magnitude of background excitatory and inhibitory firing is lowered, the gain onto a target neuron increases. This gain effect arises from two separate processes: 1) A slight hyper-polarization driven by the difference in excitatory versus inhibitory synaptic decay times, and; 2) An increase in the variance of the membrane potential (Chance et al., 2002).

Oscillatory modulation in visual cortex is traditionally modeled as either an excitatory *or* an inhibitory process. However this excitatory *or* inhibitory arrangement stands in contrast to our current knowledge of cortical connectivity and dynamics where fast and balanced EI interactions are the norm (Haider, 2006). Recent cortical micro-connectivity studies further suggest that balanced EI connectivity exists not only within cortical layers, but also between layers and across cortical regions (Markram et al., 2015; Pluta et al., 2015).

Beyond the assumption of global EI balance (Brozović et al., 2008; Haider, 2006; Litwin-Kumar et al., 2011), we assume:

1. The LIF model is sufficient to capture the effect of background noise on neural tuning (Abbott and Chance, 2005; Ayaz and Chance, 2009; Chance et al., 2002). However we note that, in modeling studies, dendritic connectivity can strongly alter cortical tuning (Behabadi et al., 2012; Jadi et al., 2012). Incorporating dendritic effects may therefore represent an important route for future work.
2. Large-scale cortical dynamics are well represented by an average firing rate model (Chaudhuri et al., 2015; David and Friston, 2003; Magri et al., 2012; Serrano et al., 2013; P. Wang and Knösche, 2013; Wong, 2006). Like the case of dendritic effects above, extending our approach to include populations of *heterogeneous* neurons represents another important future extension (Jirsa and Stefanescu, 2010; Stefanescu and Jirsa, 2008).
3. A two-population model of only excitatory and inhibitory neurons is sufficient to capture the fundamental effects of gain modulation in cortex (P. Wang and Knösche, 2013; Wilson and Cowan, 1973). Rate models with a larger numbers of populations offer a richer set of potential dynamics (Cona et al., 2014; Jirsa and Stefanescu, 2010; Sotero et al., 2007; Ursino et al., 2010), but should not alter the fundamentally biophysical EI effect on which our model relies (Chance et al., 2002).
4. The driver/modulation dichotomy (Abbott and Chance, 2005; Buzsáki and Mizuseki, 2014).
5. As a corollary of *1*, we assume single neuron properties can be generalized to predict the tuning of many homogeneous neurons in a rate model (Shriki et al., 2003; P. Wang and Knösche, 2013; Wilson and Cowan, 1973; Zandt et al., 2014).
6. Alpha oscillations are well approximated by periodic pauses, *i.e.*, as periods of below-baseline firing (Jensen and Mazaheri, 2010; Mazaheri, 2010; Vijayan and Kopell, 2012),(Bollimunta et al., 2008).

### Single neuron model

Consistent with prior experiments, oscillatory modulations were first examined in a single LIF neuron (Eq. 1-4).

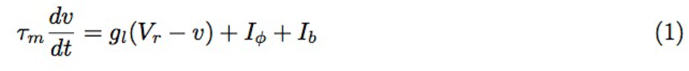

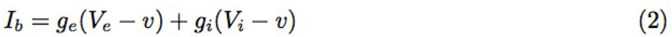

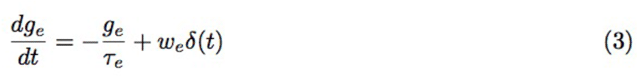

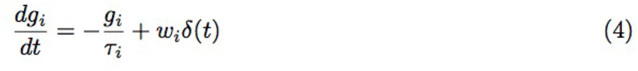

Where *v* is the membrane voltage, and *V* are the rest and Nernst reversal potentials for the neuron (*Vr*) and its synapses (*Ve*, *Vi*) and (*we*, *wi*) are the synaptic weights. *δ*(*t*) is Dirac function centered at spike time *t*. *Iφ* is the input current (see below). Following the typical LIF definition, if the membrane voltage *v* exceeded the fixed threshold *Vt*, *v* was immediately reset to *Vr* and an action potential was generated. See Table 1.

**Table 1:** Model parameters

| Parameter | Value | Units |
| --- | --- | --- |
| $\tau_m$ | 20 | milliseconds |
| $g_l$ | 10 | nanosiemens |
| $V_{reset}$ | -60 | millivolts |
| $V_r$ | -65 | millivolts |
| $V_e$ | 0 | volts |
| $V_i$ | -80 | millivolts |
| $w_e$ | 4 | nanosiemens |
| $w_i$ | 16 | nanosiemens |
| $\tau_e$ | 5 | milliseconds |
| $\tau_i$ | 10 | milliseconds |
| $b_e$ | 30-135 | Hz |
| $b_i$ | 30-135 | Hz |
| $b_0$ | 30 | Hz |
| $f$ | 10 | Hz |
| $\theta^e$ | 8 | Hz |
| $\theta^i$ | 20 | Hz |
| $w^{e,e}$ | 0.9 | - |
| $w^{i,e}$ | 1 | - |
| $w^{thal,e}$ | 12 | - |
| $w^{e,i}$ | 1 | - |
| $w^{i,i}$ | 0 | - |
| $I_\phi$ | 0-50 | millivolts |
| $S_0$ | 30 | Hz |
| $\sigma$ | 0.04 | Hz |
| $\tau_o$ | 5 | milliseconds |

### *φ*-curve

The F-I curve (*φ*) represents the relation between an input to single neuron and its firing rate output (Fig. 1a-c). We study F-I curves of neurons that are subject to a set of balanced background parameters, Ω = *{be, bi, we, wi, τe, τi}*, which act to modulate the gain of the neuron (Chance et al., 2002). Specifically, for a given *i*th neuron 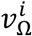, the F-I curve is the range of firing rates produced by a range of input currents, *Iφ* (Table 1).

### Background activity

Three modes of background modulation were explored. In each, the balanced condition was always maintained (see *Introduction*). In each condition only the rates were modulated, *i.e.*, *{be, bi}* ⊂ Ω. In the constant mode, the LIF neuron was subject to a fixed rate of background activity level *b*. In the oscillatory mode, the background firing rates varied with time according to a truncated cosine (Eq. 5). This rectified wave more closely resembled the synchronous pulses of activity often found in recordings of thalamic oscillations. The third mode, bursting, is a mix of the previous mode. Firing is constant at *b* until a preselected time *tf*, after which there are *k* cycles of oscillation (Eq. 5), followed by a return to *b*.

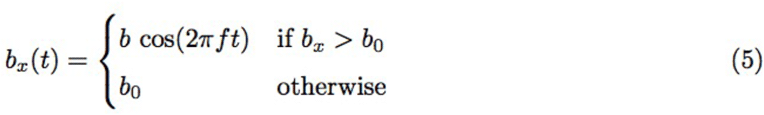

### Multi-scale rate model

Our simplified model of layer 4 in visual cortex consisted of two reciprocally coupled EI populations, as well as two thalamic inputs (Fig. 3a). One thalamic connection acted as a driver of excitatory activity, *i.e.*, having a weight that is strong (4-8x) compared to the internal connectivity of the layer model. The second input modulates the gain (output nonlinearity) of the network (discussed below).

Our formulation for population firing rate was based on the classic model of Wong and Wang (Wong, 2006), but was modified to retain nonlinear connectivity between excitatory and inhibitory populations, and neglecting NDMA synapses. The rate equations for population *x* are:

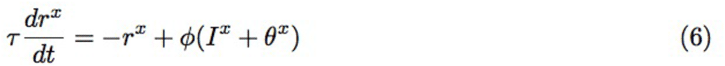

Where *x* denotes either excitatory (*e*) or inhibitory (*i*) activity, and *I*^*x*^ is the total synaptic input into the model, defined by Eqs. 7 and 8. The bias term *θ* was adjusted so the baseline firing rate the populations was approximately 6 and 12 Hz for *e* and *i* respectively (Table 1).

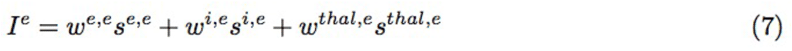

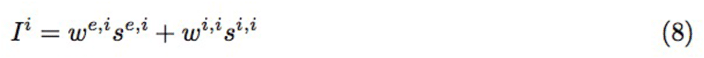

Each of the synapses *s*^*x,y*^ is modeled as single exponential function increasing or decreasing additively with the population rate *r*^*x*^ (Eq. 9). The time constant *τx,y* is either *taue* or *taui* depending on the population of *x*.

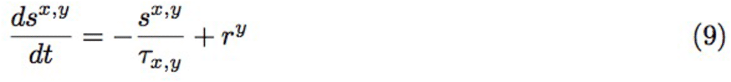

### Dynamic gain control

Altering the output nonlinearity *φ* in a rate model simulates gain modulation. The output nonlinearity in Wong and Wang (Wong, 2006) is based on a closed-form solution for balanced gain modulation taken from Abbott and Chance (Abbott and Chance, 2005). This solution was based on a simulated LIF neuron’s F-I curve as it was subjected to a range of balanced background modulations (see Fig. 1c). As in most rate models, however, this nonlinearity is static in time.

Introducing a dynamic output nonlinearity, and therefore dynamic gain control, required moving from the closed-form solution to a multi-scale simulation approach. This approached joined together a single neuron (LIF) simulation with the population rate model, with the former used to estimate the gain/output nonlinearity of the latter.

For every population time-step *dt* in Eq.6, a new instantaneous F-I curve was estimated using the single neuron model (Eqs. 1-4). That is, at every time *t* a background instantaneous parameter set Ω and an input range *I_φ_*, generating a single-unit tuning curve. This curve then defined the output nonlinearity *φ*_Ω_, subject to linear interpolation, as necessary, to account for the discrete sampling of *I_φ_*.

Substituting in the multi-scale nonlinearity give us the final rate equation (Eq. 10) with an additional constraint (Eq. 11) that keeps *I* in the range supported by *φ*_Ω_.

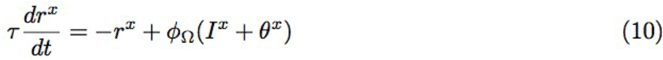

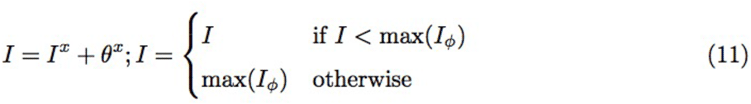

Multi-scale simulations are computationally demanding. To make ours tractable we restricted changes in *{be, bi}* to 1 Hz increments. Restricting *{be, bi}* worked in concert with a caching system that stored prior *φ*_Ω_ curves to disk. Initial “burn in” runs took 4-24 hours, while subsequent runs using caching parameters generally completed within 2-3 seconds.

### Simulated task design

#### Visual target detection

In a typical visual detection task a faint flash or grating is presented on screen for a brief period (10-20 ms). Detection of the stimulus is indicated by a button press. To simulate this task, a brief stimulus (11.7 ms) was injected into the model’s excitatory population 400 ms after trial onset. This stimulus was modeled as a 50-150 Hz burst of Poisson firing. The intensity of the stimulus was adjusted so it was detectable on 70% of trials. Detection required the excitatory population’s firing rate to exceed 2 standard deviations above the averaged pre-stimulus rate (350 to 100 ms before onset). The detection window spanned 400-420 ms following trial onset. Activity was averaged over this period during detection. Such a narrow window would be infeasible for human participants, however the stimulus onset, and neural response time-constant, are precisely known in our simulated experiment; increasing the integration windows only serves to linearly decrease *d’*.

During half the trials no stimulus was presented to the model. The combined stimulus/no stimulus trials allowed for calculation of *d’* for each simulation experiment by controlling for “false alarms” (outlined below). Each experiment contained 720 trials: 360 with stimulus presentation and 360 without.

Four experiments were performed using this task. Each manipulated only the parameters for the alpha rhythm.

*Experiment 1* compared *d’* between a constant gain condition and oscillatory gain condition, where the phase of oscillation was randomly sampled from a uniform distribution, *-*2*π,* 2*π* (Fig. 3b-e).

*Experiment 2* compared *d’* as function of oscillatory power, spanning a range of 1-3x over the default rate of 135 Hz (Fig. 3f-g).

*Experiment 3* phase-locked a 2-cycle burst of oscillation to stimulus onset, such that the stimulus was appeared in the center of trough; the highest gain period (Fig. 3j-l).

*Experiment 4* combined two V1 populations acting as separate “hemispheres” in an ipsi/contralateral visual detection task. This modified version of the model and task simulates a visual field detection task where the participant must indicate which side of the visual field a stimulus appeared. The manipulation condition in this task was ipsi-or contralateral alpha power (Fig. 3h,i).

#### Naturalistic vision

Naturalistic viewing experiments, in both humans and non-human animals, typically have the participants to watch a video recoding of natural scenes. The pattern of early visual activity under these conditions is often reported to correspond to an Ornstein–Uhlenbeck process. In such tasks, alpha activity is reported to track both stimulus intensity and attentional allocation. We therefore modeled to what degree thalamic alpha gain modulation could increase information flow in layer 4 of our modeled V1. To model natural viewing condition, a time varying stimulus *S* was sampled from an Ornstein–Uhlenbeck process and fed into the excitatory population (Eq. 12; *η* is the normal distribution with mean 0, variance 1; Fig. 4a).

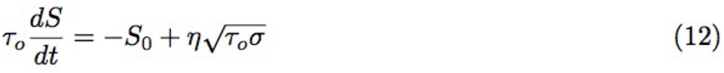

*Experiment 1* compared mutual information between the incoming naturalistic stimulus and the excitatory population’s dynamics.

*Experiment 2* was similar to *1* but featuring two V1 circuits. Here both circuits were subjected to alpha modulation, but the phase between the oscillations varied with each trial (Fig. 4f-h). Phase differences from 0 to *π* radians were explored. Mutual information was measured as in *1*.

### Analysis

#### Mutual information

Information content was estimated between the stimulus and the excitatory populations in 1 ms intervals. To allow for reliable calculation of the conditional probabilities necessary for information theoretic calculations, the summed rates were discretized into 10 integer levels. Integer levels between 4 and 30 were considered, but did not alter the patterns of results we report here. Using this new activity “alphabet”, entropy and mutual information (MI) were subsequently calculated with the pyentropy library (Ince, 2009), using the Panzeri–Treves method (Panzeri and Treves, 1996) of correcting for downward bias in estimating entropy *H* introduced by finite sampling of each time window.

#### EEG

EEG data was collected from 29 participants performing a visual target detection task (detailed below). All subjects gave informed consent approved by the UC San Diego Committee on Human Research Protections. Scalp EEG data were collected using a 64 electrode BrainVision EEG system (Brain Products) with electrode impedances kept below 10 kΩ. Data were sampled at 5000 Hz and with online reference of FCz, and later downsampled to 500 Hz.

Each participant completed at least one resting state (eyes closed) EEG recording at the beginning of the experiment. After the resting state blocks, participants performed a visual detection task. Participants sat facing a CRT monitor, with a 150 Hz refresh rate. Participants were instructed to keep their eyes fixated on a white fixation cross at the center of an otherwise black screen. At the start of the trial, horizontal lines appeared, one across the top of the screen and another across the bottom. During the time when the white lines were present the participant was instructed to covertly attend to the region between the top and bottom line and report (by pressing a button on a keyboard) if they detected a brief (13 ms) flash of light that may appear. In 80% of trials, a flash was presented, with the remaining 20% serving as catch trials, in which no stimulus was presented. Participants behaviour was thresholded to 50% detection rate, by adjusting the luminance of the presented stimuli.

Task data comes from two variants of the visual detection task outlined above. In the first version, 15 subjects (age 19-28, 8 female) responded if they detected a flash of light at one of two peripheral locations, which could occur between 1000 - 2500 ms post cue on set, with stimulus presentation implemented in Psychopy(Peirce, 2007), running in Python. A further 14 subjects (age 18-21, 10 female) performed a variant of the task in which 4 possible peripheral locations were used, with flashes presented randomly between 500 and 1500 ms post cue onset. In this variant, stimuli were presented with the Psychtoolbox-3 presentation toolbox (Brainard, 1997) running on Matlab (MATLAB R2014a, The Mathworks).

Data used for this investigation comprised the time-series data for the first two minutes of continuous recordings from the first resting state block for each participant, as well as the continuous data for the first two minutes of the first test block (post behavioral thresholding) for each participant, from a single posterior electrode (Oz). To estimate alpha power, the instantaneous power in the 8-12 Hz range data from extracted by taking the absolute values of the Hilbert transform which was subjected to a burst detection procedure (below).

#### ECoG

These data were previously published (Coon et al., 2016), where complete acquisition and processing details can be obtained. Participants gave informed consent for the study, which was approved by the Institutional Review Board of Albany Medical College and the Human Research Protections Office of the U.S. Army Medical Research and Materiel Command. In brief, ECoG data was collected from four participants using eight 16-channel USBamp biosignal acquisition devices (g.tec, Graz, Austria). Of these, in our analyses two patients showed strong alpha in their more posterior electrodes. Data was digitized at 1200 Hz and visually inspected offline; channels that did not contain clear ECoG signals were removed prior to analysis. Data was acquired while patients completed a modified Posner cueing task (Posner, 1980), however our analysis did not examine task-specific effects and instead focused on the general relationship between alpha and gamma power, as our model’s predictions are not contingent on a set task-state.

#### Burst detection

Periods of oscillations (*i.e.*, bursts) were detected based on a modified version of the method wherein they detected bursts by comparing instantaneous power to median power in the channel of interest (Feingold et al., 2015). However for very stable oscillations and very intermittent oscillations, this criterion becomes unreliable. To detect bursts we instead compared power in the alpha band (8-12 Hz) to power in the 1/*f* background of the power spectrum. An instantaneous estimate of background power was found by averaging power from two narrowband filters on either side of the alpha band—3-5 and 12-14 Hz. A similar approach was validated for human alpha (Whitten et al., 2011), but calculating full power spectra at every time-step. Using our approach, a burst was then said to begin whenever the alpha (8-12 Hz) power exceeded 2 times background power, and ended when power dropped below 1.5 times background power. Burst length is defined by (*t*offset-*t*onset).

## ACKNOWLEDGEMENTS

We thank Tom Donoghue as well as William Coon and Gerwin Schalk for graciously sharing EEG and ECoG data, respectively. This work was supported by a Sloan Research Fellowship, the National Science Foundation (1736028), and the University of California, San Diego (UCSD) Qualcomm Institute, California Institute for Telecommunications and Information sTechnology, Strategic Research Opportunities Program.

## AUTHOR CONTRIBUTIONS

E.J.P. and B.V. conceived of the research and approach, analyzed the human electrophysiological data, and wrote the manuscript. E.J.P. designed, implemented, and analyzed the computational models. The authors declare no competing financial interests.

